# Determination of permissive and restraining cancer-associated fibroblast (DeCAF) subtypes

**DOI:** 10.1101/2024.05.14.594197

**Authors:** Xianlu Laura Peng, Elena V. Kharitonova, Yi Xu, Joseph F. Kearney, Changfei Luan, Priscilla S. Chan, Arthi Hariharan, Ian C. McCabe, John R. Leary, Ashley B. Morrison, Hannah E. Trembath, Michelle E. LaBella, Silvia G. Herera Loeza, Ashley Cliff, Hong Jin Kim, Brian A. Belt, Roheena Z. Panni, David C. Linehan, Jeffrey S Damrauer, Alina C. Iuga, William Y. Kim, Naim U. Rashid, Jen Jen Yeh

## Abstract

Cancer-associated fibroblast (CAF) subpopulations in pancreatic ductal adenocarcinoma (PDAC) have been identified using single-cell RNA sequencing (scRNAseq) with divergent characteristics, but their clinical relevance remains unclear. We translate scRNAseq-derived CAF cell-subpopulation-specific marker genes to bulk RNAseq data, and develop a single-sample classifier, DeCAF, for the classification of clinically <u>rest</u>raining and <u>perm</u>issive CAF subtypes. We validate DeCAF in 19 independent bulk transcriptomic datasets across four tumor types (PDAC, mesothelioma, bladder and renal cell carcinoma). DeCAF subtypes have distinct histology features, immune landscapes, and are prognostic and predict response to therapy across cancer types. We demonstrate that DeCAF is clinically replicable and robust for the classification of CAF subtypes in patients for multiple tumor types, providing a better framework for the future development and translation of therapies against permissive CAF subtypes and preservation of restraining CAF subtypes.

**Significance:** We introduce a replicable and robust classifier, DeCAF, that delineates the significance of the role of permissive and restraining CAF subtypes in cancer patients. DeCAF is clinically tractable, prognostic and predictive of treatment response in multiple cancer types and lays the translational groundwork for the preclinical and clinical development of CAF subtype specific therapies.

## INTRODUCTION

It is widely recognized that the PDAC tumor microenvironment (TME) plays an important role, and can be both tumor restraining and tumor permissive^1–4^. PDAC is characterized by an extremely dense desmoplasia, represented as a complex mixture of extracellular matrix (ECM), blood vessels, immune cells, as well as cancer-associated fibroblasts (CAF)^5–10^ . CAFs are key regulators in the TME. Clinical trials attempting to target the PDAC TME have been disappointing^11–14^. These results may be partially explained by the loss of tumor restraint with genetic depletion of CAFs in genetically engineered mouse models^2,3,15^.

We previously reported two PDAC stroma groups, “activated” and “normal”, where patients with “activated stroma” had decreased survival relative to normal stroma^16^. Maurer et al. used microdissected patient samples to derive two TME groups called “ECM-rich” and “immune-rich” stroma, with “ECM-rich” showing shorter survival^17^. With recent advances in single cell RNA sequencing (scRNAseq) technology, studies on the PDAC stroma have rapidly shifted to the study of individual CAF and immune cell populations. In a landmark study, Elyada et. al identified “myCAF” and “iCAF” cell clusters described initially in preclinical murine studies and validated in a human scRNAseq dataset (Elyada-sc) that enriched for CAF cells using fluorescence-activated cell sorting (FACS)^18,19^. More recently, additional CAF subpopulations (e.g. “csCAF”, “meCAF”, etc.) have been described^20–22^, as well as CAF subtypes with differential histology features^23,24^, and CAF related gene programs^25^. However, the clinical significance of previously described CAF subpopulations remains unproven, despite interest in targeting iCAFs due to their tumor promoting behavior in preclinical studies.

Using SCISSORS, a method that we previously developed to sensitively cluster rare cells and identify highly cell-subpopulation-specific marker genes in scRNAseq, we identify uniquely expressed marker genes for CAF clusters that robustly translate to bulk RNAseq^26^. We show that these marker genes are specific to fibroblasts and are not confounded by expression in immune cells. We successfully translate these CAF marker genes to develop and validate a single sample classifier (SSC), DeCAF (<u>De</u>termination of permissive and restraining <u>c</u>ancer- <u>a</u>ssociated fibroblast subtypes), that robustly and replicably classifies <u>perm</u>issive (permCAF) and <u>rest</u>raining CAF (restCAF) subtypes in 19 independent bulk transcriptomic datasets of patient samples across four tumor types. We find that DeCAF subtypes are independently prognostic and associated with distinct histologic features. Furthermore, DeCAF subtype tumors are associated with different immune landscapes and are associated with treatment response in a PDAC phase Ib trial of FOLFIRINOX and CCR2 inhibition^27^. In mesothelioma, urothelial and clear cell renal cell carcinoma (cRCC), DeCAF subtypes are prognostic and in urothelial and cRCC associated with response to anti-PD-L1 therapy. Here, we show that DeCAF subtypes are robust, replicable and elucidate the relationship between CAF subtypes, prognosis, and treatment response in patients with mesothelioma, cRCC, bladder and pancreatic cancers.

## RESULTS

### Development and external validation of DeCAF

The clinical relevance of CAF subpopulations derived from scRNAseq remains unclear largely due to the paucity of samples scRNAseq from patients. Therefore, we set out to translate exemplar or marker genes derived from scRNAseq for use in bulk transcriptomics data. We previously showed that the basal-like and classical cell-subpopulation-specific marker genes derived from scRNAseq may be translated to cluster bulk RNAseq of PDAC samples, and highly recapitulated the clinically validated PDAC tumor subtypes^26^. These marker genes were identified using a previously developed tool, SCISSORS, which sensitively identifies rare cell subpopulations as low as 0.092% of the cell population and accurately identifies marker genes of high specificity. Because of this, we hypothesized that the CAF cell-subpopulation-specific marker genes derived by SCISSORS may also be translated to bulk RNAseq for the classification of CAF subtypes in patient samples (Supplementary Table 1). First, we evaluated 11 publicly available bulk RNAseq and microarray datasets containing 1,432 primary PDAC patients using consensus clustering (CC) (Methods, Supplementary Table 2, 3). In all datasets, the SCISSORS CAF genes separate patient samples into two clusters (Supplementary Fig. 1). In a meta-analysis, we found two divergent clusters: one patient cluster had tumor <u>perm</u>issive CAF (permCAF) with a significantly shorter median OS (mOS) of 20.33 months (mos), while the other patient cluster had tumor restraining CAF (restCAF) with a longer mOS of 30.19 mos (p = 0.0036, HR = 1.40 [95% CI 1.317-1.725], Supplementary Table 4). The naming of permCAF and restCAF describes the clinical implications of our results and avoids conflict with the existing preclinical nomenclature.

However, clustering techniques have many real-world limitations, such as their inability to assign subtypes to individual patients and the lack of stability in existing subtype assignments due to the re-clustering required for the addition of new samples to existing data and can be inherently deceptive when evaluating samples of limited numbers and diversity. Thus, we developed a robust SSC, DeCAF (Fig. 1a, Supplementary Table 5), to predict permCAF and restCAF subtypes in individual patients, using CC based training labels derived in four large bulk gene expression datasets (Methods, training datasets: CPTAC, Dijk, Moffitt GSE71729, TCGA PAAD, Supplementary Table 2, 3, Supplementary Fig. 2). A key element of our method includes the utilization of our CAF marker genes derived by SCISSORS to avoid the confounding of the expression of these genes in tumor and other tissue types such as immune cells. The final DeCAF classifier uses rank-derived predictors through the k Top Scoring Pair (kTSP) (https://github.com/jjyeh-unc/decaf) (Methods); the same approach that we employed for the PurIST PDAC tumor classifier^28^. This approach avoids using raw expression values which reduced its dependence on between sample/study normalization, simplifying data integration over different studies^28–31^ and during prediction on new samples.

**Fig. 1:**
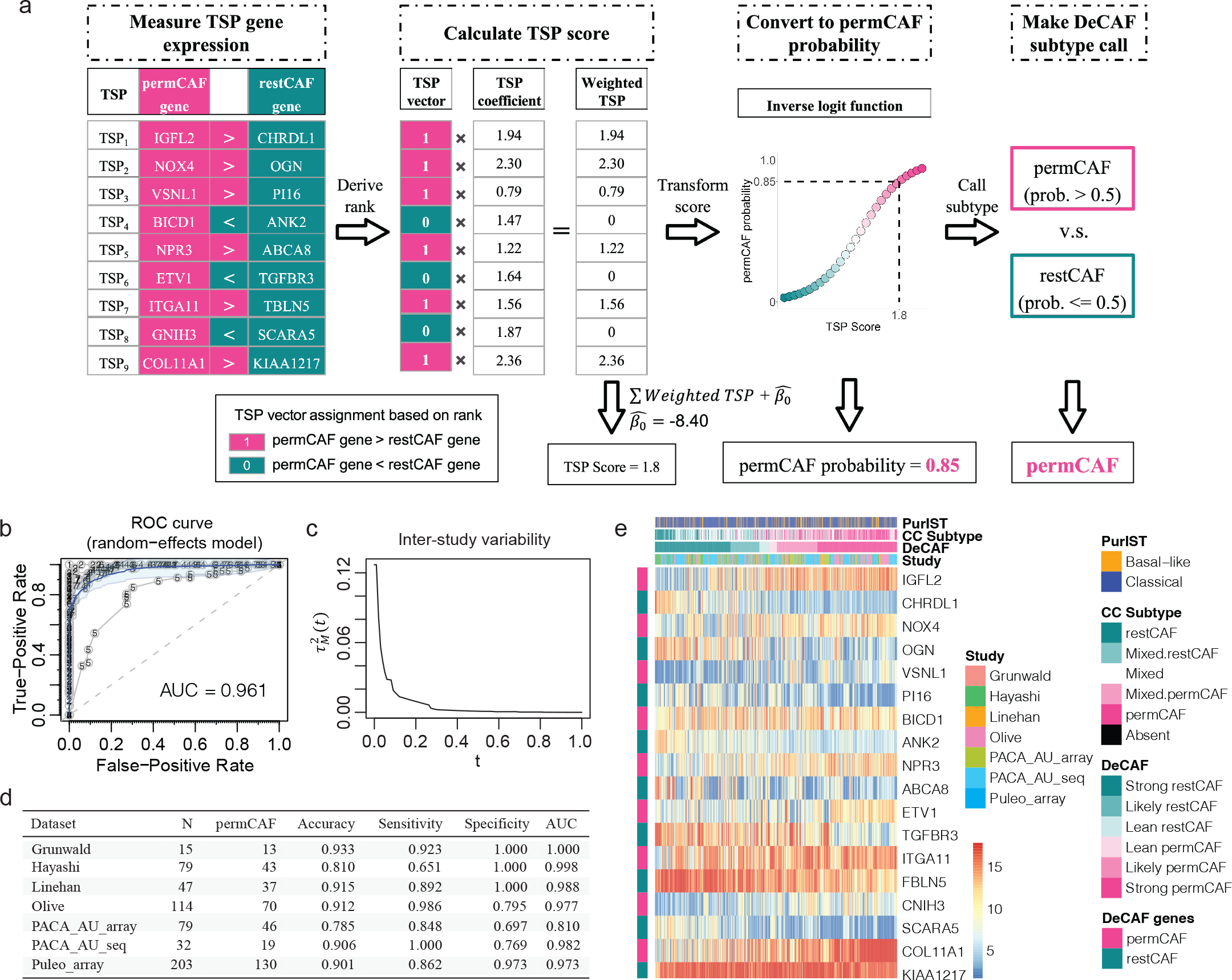
Development and external validation of DeCAF. a, Framework of DeCAF, using 9 pairs of TSP genes to derive a DeCAF permCAF probability which is then converted to the final permCAF vs restCAF call for a single sample. **b,** ROC curve for the evaluation of DeCAF calls against gold standard labels, which were made by consensus clustering (CC) using SCISSORS CAF genes. **c,** Inter-study variability for the evaluation of DeCAF calls. **d,** Summary statistics of the evaluation of DeCAF subtype calls in each validation dataset. **e,** Heatmap showing SCISSORS-CC calls and DeCAF subtype calls in the pooled samples.

To assess the quality of our prediction model, we first evaluate the cross-validation error of the final model in our Training Group samples. We find that the internal leave-one-out cross validation error for DeCAF in the Training Group is low (4.0%). To evaluate the overall classification performance of DeCAF across additional studies, we compare the DeCAF subtype calls to the SCISSORS subtype calls in an independent validation dataset of seven patient cohorts (Supplementary Table 2, 3). First, we applied a nonparametric meta-analysis approach to obtain a consensus ROC curve based on the individual ROC curves from each validation study. We found that the overall consensus Area Under the Curve (AUC) is high, with a value of 0.961 (Fig. 1b). The estimated interstudy variability of these ROC curves with respect to predicted permCAF score threshold *t* is very low at our standard threshold of *t* = 0.5 or greater (Fig. 1c). Furthermore, sensitivities and specificities were often high at this threshold, and AUC values were similarly strong (> 0.8) (Fig. 1d). Across validation datasets, we find that the pooled samples strongly segregate by CC subtype when sorted by their DeCAF score (i.e. permCAF probability), despite diverse studies of origin (Fig. 1e). This suggests that our methodology avoids potential study-level effects. As expected, the relative expression of classifier genes within each classifier TSP (paired rows) strongly discriminates between subtypes in each sample, forming the basis of our robust TSP-oriented approach for subtype prediction (Fig. 1e). Our results support that DeCAF is robust and replicable, supporting our comprehensive investigation to evaluate the biological, pathological and clinical relevance of permCAF vs restCAF in patient samples.

### Cell specificity of DeCAF in scRNAseq

Elyada et al. has previously described two PDAC CAF subtypes, namely myCAF and iCAF^18,19^. Using the scRNAseq dataset generated by Elyada et al. (Elyada-sc), we found that within the fibroblast cell clusters, our permCAF genes showed overlapping expressions with Elyada myCAF genes, and our restCAF genes showed overlapping expressions with Elyada iCAF genes (Fig. 2a,b). However, we found that our DeCAF and SCISSORS genes are uniquely expressed within only CAF cells and not confounded by other cell types, e.g. epithelial cells and immune cells, demonstrating the high specificity of our CAF marker genes (Fig. 2b). In addition, DeCAF permCAF and restCAF genes were distinctively expressed by different clusters of CAF subpopulations, illustrating the high CAF cell-subpopulation specificity of the genes (Fig. 2c). As expected, DeCAF and SCISSORS genes showed very limited overlap with the CAF marker genes identified in other scRNAseq-based studies^16–18,20–25^ (Supplementary Fig. 3).

**Fig. 2:**
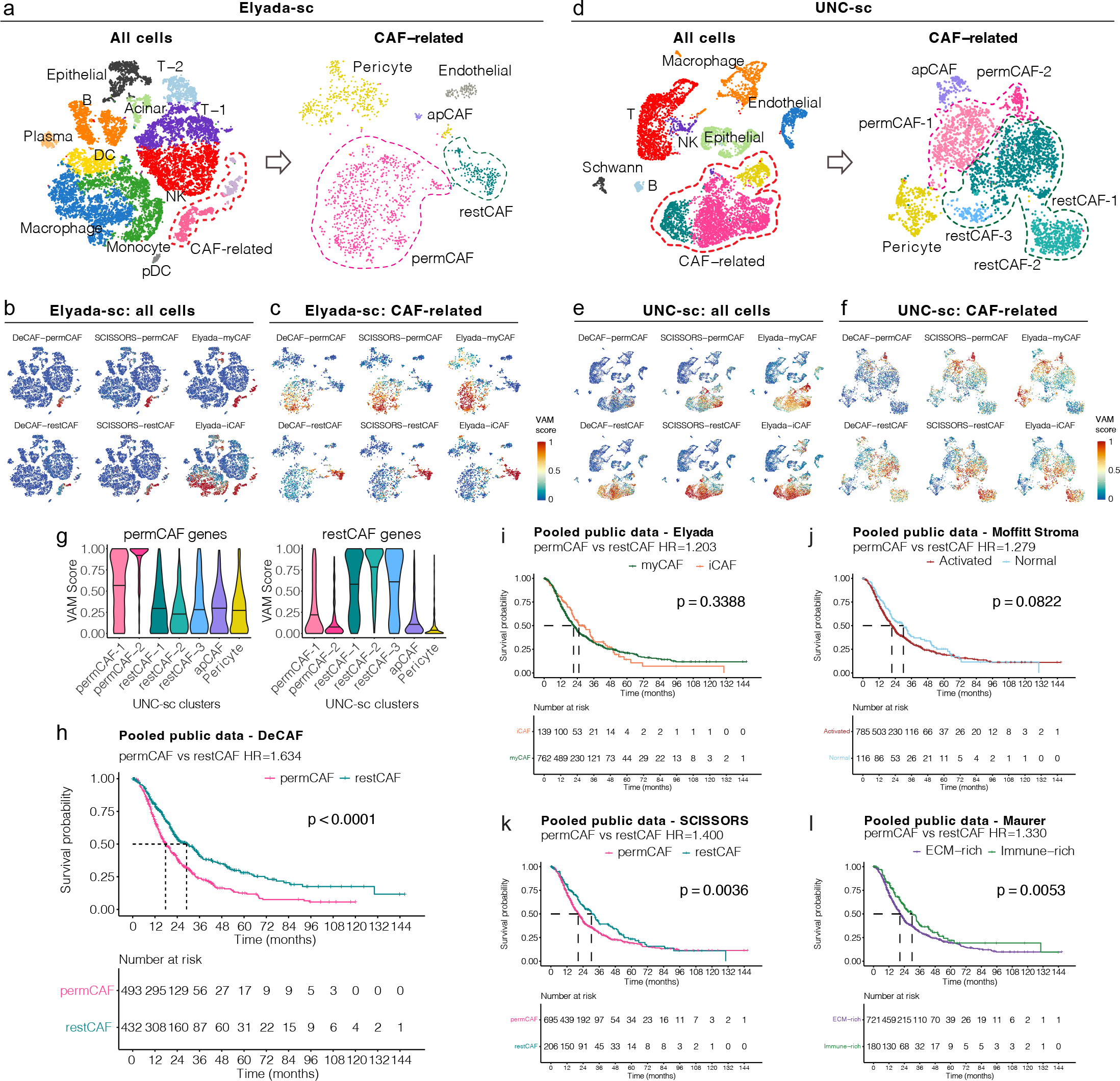
Benchmarking DeCAF with existing CAF subtypes. a&d, Clustering of CAF cells by applying SCISSORS on the Elyada-sc and UNC-sc scRNAseq datasets respectively. **b&e,** Gene set enrichment VAM scores for the DeCAF, SCISSORS and Elyada CAF genes in each cell shown on the UMAP of all cells in Elyada-sc and UNC-sc dataset. **c&f,** Gene set enrichment VAM scores for the DeCAF, SCISSORS and Elyada CAF genes in each cell shown on the UMAP of CAF-related cells in Elyada-sc and UNC-sc dataset. **g,** Violin plots showing gene set enrichment VAM scores, of SCISSORS marker genes in UNC-sc CAF-related cells. **h-l,** Kaplan-Meier plots showing patient overall survival (OS) of CAF subtypes called by DeCAF, as well as CC-based methods using SCISSORS, Elyada, Moffitt stroma and Maurer CAF genes. Stratified log-rank test.

To further validate the DeCAF genes, we evaluated an independent scRNAseq dataset with six primary PDAC samples (UNC-sc, Supplementary Table 6). Analyzing the UNC-sc data by SCISSORS, we identified fibroblast cells, along with other known cell types in PDAC samples, including epithelial cells, endothelial cells, as well as different types of immune cells (Fig. 2d). Here, we again found that the DeCAF genes are more uniquely expressed within fibroblast cells, compared to Elyada genes (Fig. 2e). As part of SCISSORS, fibroblast cells are then reclustered, where we find that permCAF and restCAF genes are distinct to different fibroblast subclusters (Fig. 2f,g). Therefore, we show that at the single-cell level, the DeCAF genes are unique and specific markers for human PDAC CAF cell subpopulations.

### Benchmarking prognostic impact of DeCAF with existing subtypes

To benchmark the prognostic impact of DeCAF against other comparable methods, we performed a meta-analysis of the pooled datasets with OS data available (Supplementary Table 4). We found that the patients with permCAF subtype tumors (mOS 17.70 mos) have significantly shorter OS than patients with restCAF subtype tumors (mOS 29.04 mos) (Fig. 2h, p < 0.0001, stratified HR = 1.634, [95% CI 1.375-1.943]). Next, we used published CAF subtyping gene signatures from Elyada et al.^18^, Moffitt et al.^16^, and Maurer et al.^17^ to derive subtype calls in each of the 11 datasets for comparison (Supplementary Table 3). In contrast to DeCAF, we found that the Elyada myCAF and iCAF gene sets were not prognostic (Fig. 2i, mOS: 21.32 vs 25.20 mos, p = 0.339, HR = 1.203 [95% CI 1.532-0.945]). The Moffitt stroma schema demonstrated shorter survival for patients with tumors with activated stroma relative to normal stroma but did not reach significance in our pooled datasets (Fig. 2j, mOS: 21.45 vs 30.0 mos, p = 0.082, HR = 1.279 [95% CI 0.991-1.651]). Similar to SCISSORS (Fig. 2k), we found that Maurer ECM-rich and Immune-rich bulk RNAseq gene signatures are associated with differences in survival (Fig. 2l, mOS: 20.53 vs 30.03 mos, p = 0.005, HR = 1.33 [95% CI 1.047- 1.69]). However, DeCAF subtypes showed the largest and most significant difference in outcome (Fig. 2h).

### DeCAF and tumor-intrinsic subtypes are independently prognostic

It is well validated that PDAC patients with basal-like and classical tumors have significantly different OS^16,28,32,33^. We hypothesize that CAF subtypes will also impact tumor behavior. First, we compared the association of DeCAF subtypes with basal-like and classical tumor-intrinsic subtypes as defined by PurIST^28^. We found that 64.5% (N=167) of basal-like tumors had a permCAF subtype, but only 35.5% of basal-like tumors had a restCAF subtype (N=92). No difference was seen in CAF subtypes within classical tumors: 49.7% permCAF (N=529) vs 50.3% restCAF (N = 536), suggesting an affinity for basal-like tumors to be a permCAF subtype (p < 0.001, Fisher’s exact test, Fig. 3a,b). Basal-like tumors also have higher permCAF probability scores, suggesting an increase permissive CAF subtype (Fig. 3c). Patients with PurIST basal-like and DeCAF permCAF subtype tumors had the shortest OS, while patients with PurIST classical and DeCAF restCAF subtype tumors had the longest OS (11.01 mos vs 30.43 mos, p < 0.001, Fig. 3d).To determine the relationship between DeCAF and PDAC tumor subtypes^16,28,32^, we performed a multivariable stratified cox proportional hazard model for the pooled public datasets including DeCAF and PurIST subtypes as variables. We found that both PurIST tumor subtype and the DeCAF subtype were independently associated with survival (p < 0.001 for both, stratified Cox proportional hazards model, Fig. 3e).

**Fig. 3:**
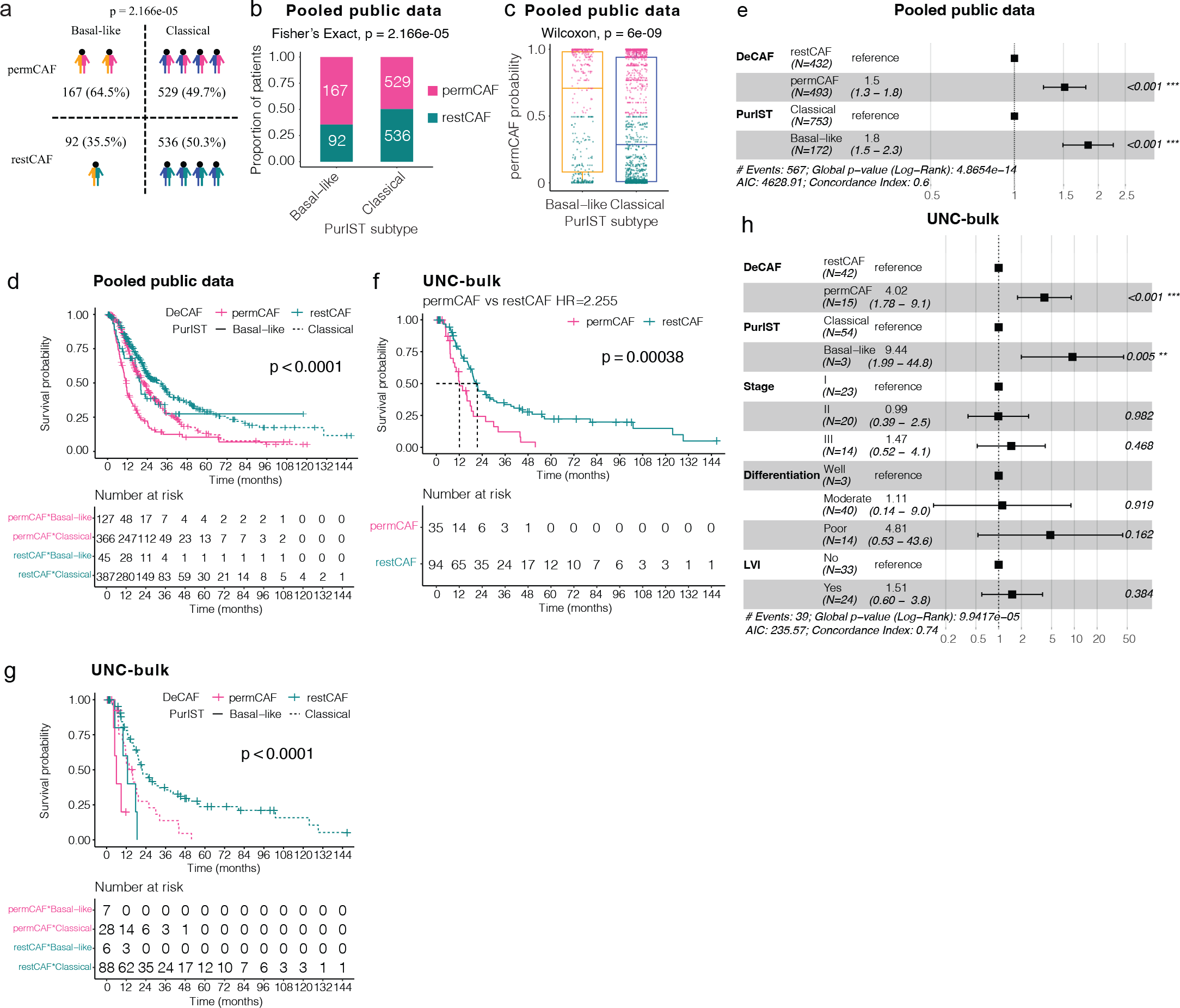
DeCAF independently predicting OS. a, Intersection of DeCAF with PurIST subtypes (p < 0.001, Fisher’s exact test). **b,** Proportion of patients in PurIST basal-like and classical tumor subtypes with permCAF or restCAF subtypes (p<0.001, Fisher’s exact test). **c,** Comparison of permCAF probability with PurIST tumor subtypes (p < 0.001, Wilcoxon rank-sum test). **d,** patients with combination of DeCAF and PurIST tumor subtypes in pooled public datasets (p < 0.001, log-rank test, stratified datasets). **e,** Multivariable Cox proportional hazards model in the pooled public datasets stratified by datasets. **f,** Kaplan-Meier plot showing OS of patients with permCAF vs restCAF subtypes (p = 0.00038, log-rank test) and **g,** of patients in the context of combined DeCAF and PurIST tumor subtypes in the UNC-bulk dataset (p < 0.0001, log-rank test). **h,** Multivariable Cox proportional hazards model in the UNC-bulk dataset.

Next, we applied DeCAF to another independent patient cohort of primary PDAC at UNC where clinical and pathology variables were available (UNC-bulk, N = 129, Supplementary Table 7) and find that DeCAF subtypes are again associated with OS (HR = 2.255, 95% CI 1.423-3.574, p < 0.001, log-rank test) (Fig. 3f). We saw similar additive effects of DeCAF and PurIST subtypes and their relationship to OS (p < 0.001, log-rank test, Fig. 3g). In an univariable analysis, we found that the restCAF subtype was associated with chronic pancreatitis on pathology (p=0.008, Fisher’s exact test), although chronic pancreatitis had no association with OS (p=0.3, Supplementary Table 8). In a multivariable analysis of patients who had complete (R0) resections, DeCAF and PurIST subtypes remained independently prognostic when including the variables stage, differentiation and lymphovascular invasion (LVI) (DeCAF p < 0.001 and PurIST p = 0.005, Cox proportional hazards model, Fig. 3h).

### Pathology differences in DeCAF subtypes

Differential pathology features of PDAC CAF subtypes have been described, where the stroma of a subgroup of patients were described as collagen-enriched^23,24^. To investigate the association of DeCAF subtypes with pathological features, hematoxylin and eosin stained slides (n = 106) were reviewed by a pathologist blinded to the subtype calls in the UNC-bulk dataset (Supplementary Table 7). Samples were annotated as either myxoid, fibrous or fibromyxoid with the fibromyxoid subtypes delineating a mixed appearance with the dominant histology noted^34^ (Fig. 4a). We found that the restCAF subtype tumors were associated with a fibrous (51/74, 68.9%) compared to a myxoid stroma histology (23/74, 31.1%) (p = 0.018, Fisher’s exact test). Samples with myxoid dominant histology showed significantly more permCAFness (i.e. higher permCAF probability) compared to fibrous histology samples (p = 0.007, Wilcoxon rank-sum test, Fig. 4b). The mixed histology type, fibromyxoid, showed intermediate permCAF probability, supporting that the DeCAF score is associated with a histologic continuum (p = 0.002, Kruskal-Wallis test, Fig. 4c). Interestingly, stroma histology alone was prognostic, with myxoid histology associated with the shortest OS, fibromyxoid with intermediate OS, and fibrous the longest (p = 0.0043, log-rank test, Fig. 4d,e). In the TCGA_PAAD dataset, as Grünwald et al. previously described subTME types based on pathology features, we sought to examine the relationship between DeCAF and the subTME types. We found that 89.6% (60/67) of permCAF subtype tumors had a reactive/intermediate subTME, compared to 10.4% (7/67) of them had a deserted subTME (Fig. 4f), while 43.5% (40/92) of restCAF subtype tumors had a reactive/intermediate subTME compared to 56.5% (52/92) that had a deserted subTME (Fig. 4f, p = 9.582e-10, Fisher’s exact test). In contrast to Grünwald et al. where patients with reactive or deserted subTME did not show prognostic differences^23^, we found that patients with fibrous stroma had longer mOS 20.6 mos compared to mOS 18.4 mos in patients with myxoid features (p = 0.004, Fig. 4d,e). However, the DeCAF subtypes show the largest and most significant prognostic separation (Fig. 2h, Fig. 3f).

**Fig. 4:**
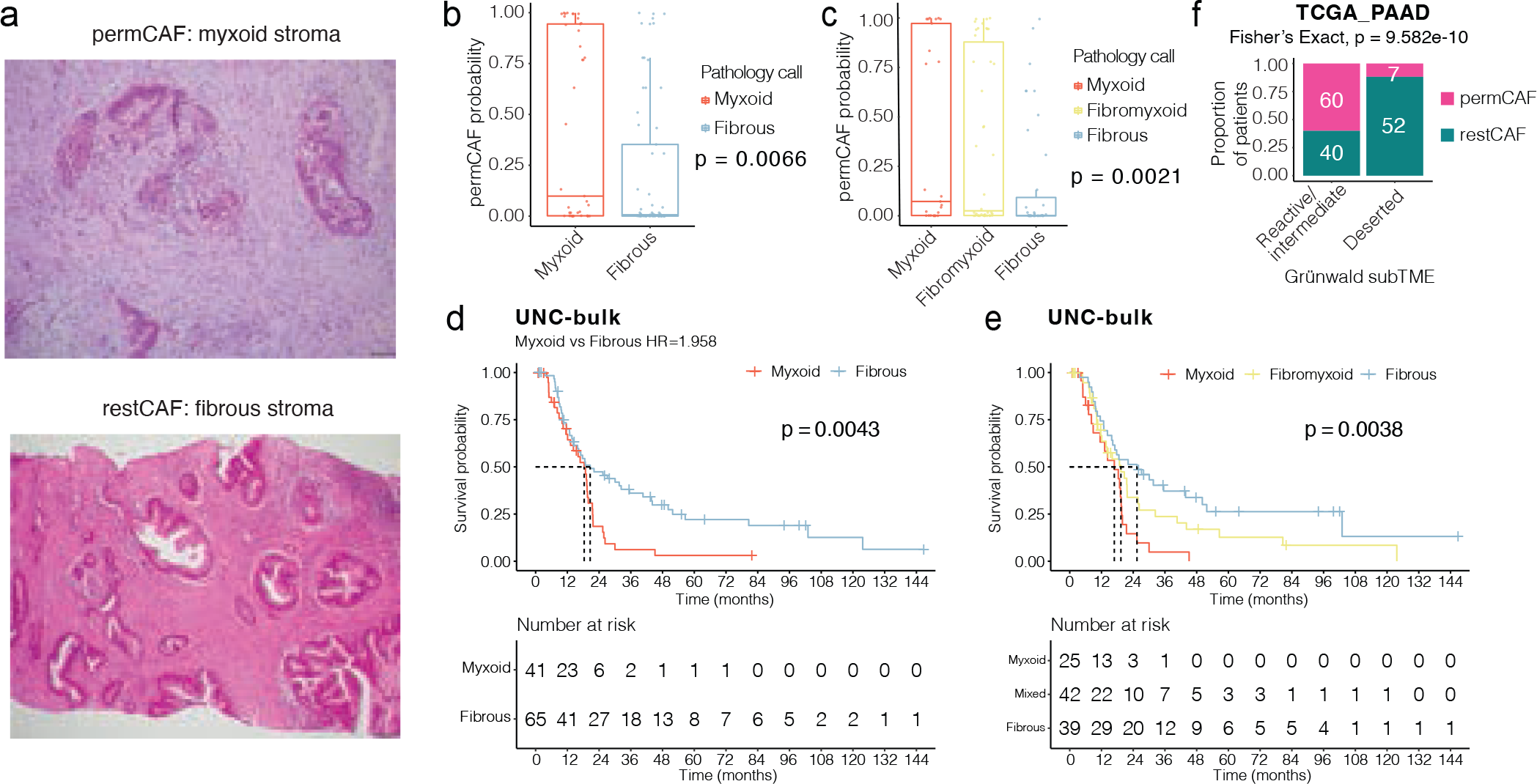
Pathology differences in DeCAF subtypes. a, Representative hematoxylin and eosin stained slides showing a permCAF subtype sample with myxoid stroma and a restCAF subtype sample with fibrous stroma. **b, c,** Boxplots comparing DeCAF probability in samples described as having a predominant myxoid vs fibrous stroma (p = 0.007, Wilcoxon rank-sum test); or having a myxoid, fibromyxoid and fibrous stroma (p = 0.002, Kruskal-Wallis test). **d, e,** Kaplan-Meier plot showing OS of patients with the different stroma histologies in the UNC-bulk dataset using a predominant classification of myxoid and fibrous stroma (p = 0.004, log-rank test), or including a fibromyxoid classification (p < 0.001, log-rank test). **f,** Proportion of patients in Grünwald reactive/intermediate and deserted tumors in TCGA_PAAD with permCAF and restCAF subtypes (p < 0.001, Fisher’s exact test).

### DeCAF subtypes in mesothelioma, cRCC and urothelial bladder carcinoma

Similarities in CAFs across cancer types have been reported^35,36^. To determine if DeCAF subtypes are seen other cancer types, we evaluated the TCGA Pan-Cancer datasets and found that DeCAF subtypes are similarly prognostic in malignant pleural mesothelioma (MESO, HR=2.056, 95% CI 1.1, 3.84, p = 0.021), cRCC (KIRC, HR=2.138, 95% CI 1.342, 3.407, p = 0.0011) and urothelial bladder (BLCA, HR 1.6, 95% CI 1, 2.6, p = 0.043) carcinomas (Fig. 5a-c).

**Fig. 5:**
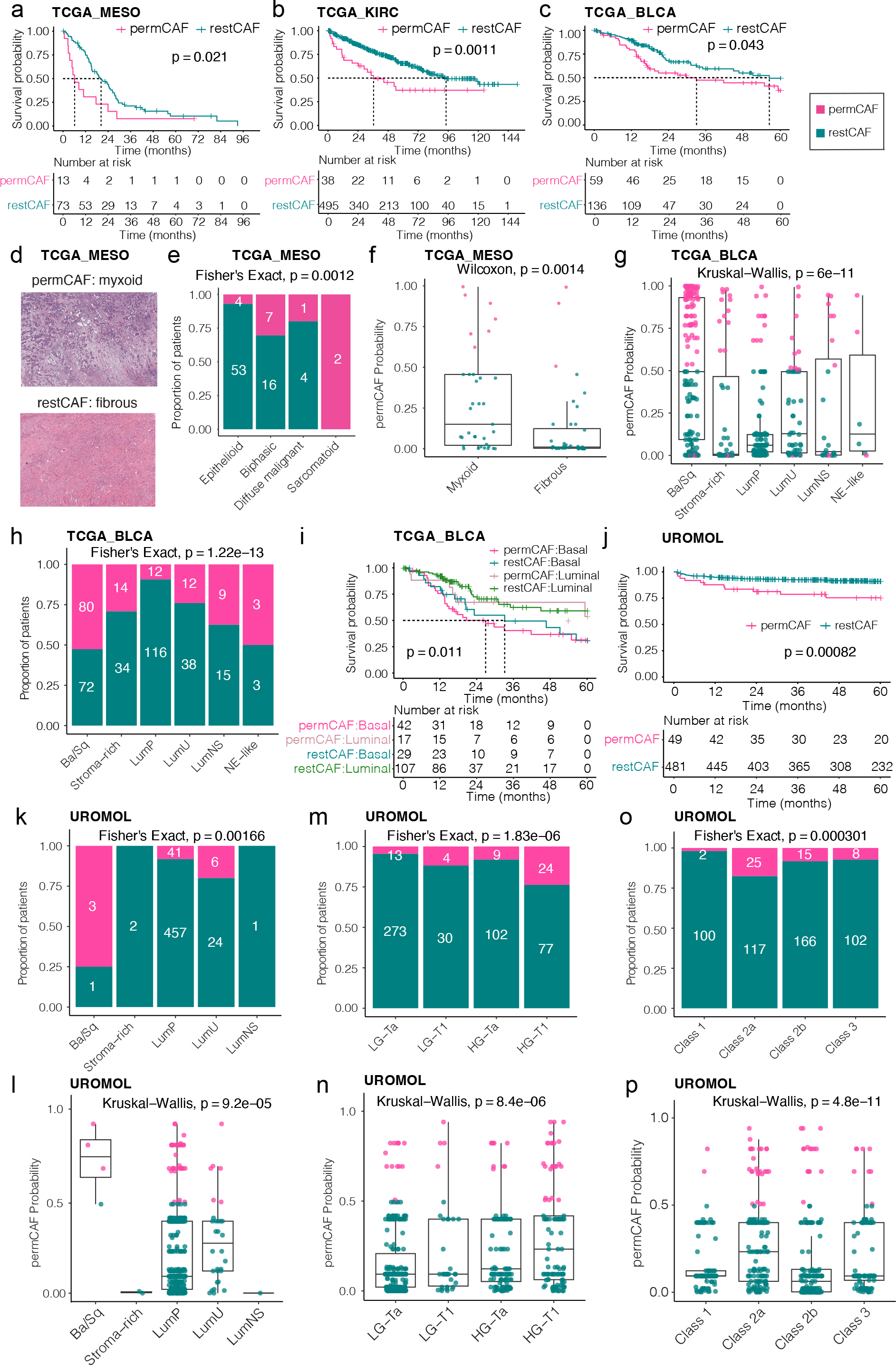

Given the clinical similarities of fibrosis that characterizes both MESO^37^ and PDAC, we hypothesized that there may also be similar pathology findings that explain the relevance of DeCAF subtypes in MESO. We found that DeCAF subtypes were associated with histological type (p = 0.001) with 93% (53/57) of epithelioid type tumors having a restCAF subtype (Fig.5d,e). Pathologist review of the stroma showed similar findings as PDAC, where a myxoid stroma was associated with higher permCAF probability (i.e. permCAFness) compared to a fibrous histology (Wilcoxon, p = 0.001, Fig.5f).

In bladder cancer where consensus subtypes have been described including a basal bladder subtype (Ba/Sq) with similar gene expression to basal-like PDAC, we found that similar to PDAC, the Ba/Sq bladder consensus subtype was enriched in the permCAF subtype, with 52.0% (80/152, p = 1.22e-13, Fisher’s exact test, Fig.5g) of Ba/Sq subtype patients having a permCAF subtype and highest permCAF probability (Fig. 5h) in the TCGA BLCA dataset. In addition, we also see a relationship between Ba/Sq tumor subtype and DeCAF subtype (p = 0.011, log-rank test, Fig. 5i). In a dataset of non-muscle invasive bladder cancer (NMIBC), UROMOL^38^, we find that patients with permCAF subtype tumors have a shorter progression free survival to developing MIBC (PFS, p=0.00082, log-rank test) (Fig. 5j). Using the BLCA consensus subtyping schema, 75% (3/4, p= 0.00166, Fisher’s exact test) of Ba/Sq tumors had a permCAF subtype and had the highest permCAF probability scores (Fig. 5k,l). Using standard clinical groupings of NMIBC, we found that permCAF prevalence and higher permCAF probability scores (i.e. permCAFness) was associated with higher grade (p = 8.7e-06, Kruskal-Wallis test, Fig. 5m,n) and most enriched in Class 2a NMIBC [18% (25/142), p = 0.000301, Fisher’s exact test], the most aggressive class of NMIBC (Fig. 5o,p).

Taken together, our results show that the presence of the permCAF subtype is associated with poor prognosis in multiple cancer types with similar histology and tumor subtype associations that we find in PDAC.

### Immune landscape of permCAF and restCAF subtype tumors

We showed that DeCAF genes have minimal expression in immune cells in two scRNAseq datasets (Fig. 2b,e). However, we hypothesize that CAF subtypes may provide different TMEs that support different immune landscapes. Therefore, we used CIBERSORT^39^ with LM22 as the reference to deconvolve the fractions of 22 types of immune cells in each of our 12 datasets (Fig. 6a, Supplementary Table 9). We found that the immune landscape was more immunosuppressive in permCAF, with an average enrichment of 1.3-fold in Tregs in 6 datasets, 1.9-fold neutrophils in 5 datasets, 1.8-fold in M0 macrophages in 10 datasets, 1.1-fold in M2 macrophages in 6 datasets (Fig. 6a). In contrast, restCAF subtype tumors had an average enrichment of 1.3-fold in M0 macrophages in 4 datasets, 1.5-fold in CD8+ T cells in 7 datasets, and 1.8-fold in naïve B cells in 6 datasets (Fig. 6a). In addition, restCAF subtype tumors showed significantly higher CD8/Treg ratios in 8 datasets (Fig. 6b), suggesting that patients with different DeCAF subtype tumors may show more favorable response to certain immunotherapies compared to patients with permCAF subtype tumors^40,41^.

**Fig 6:**
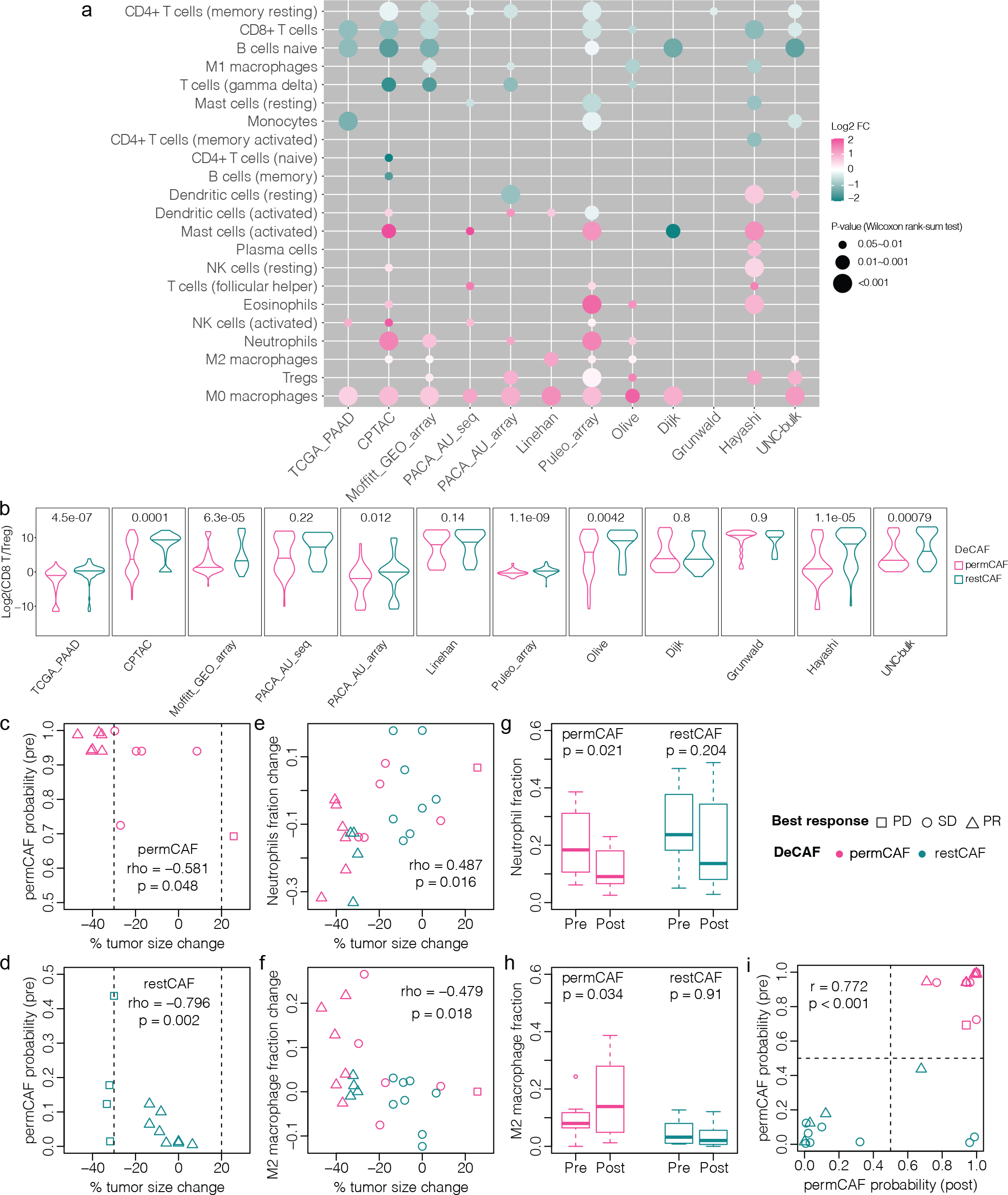
**Immune landscape of permCAF and restCAF subtype PDAC tumors. a**, Comparison of immune cell fractions deconvolved using CIBERSORT (reference: LM22) across 12 datasets. The color of the dots represents the log2 fold change (FC) of the fractions between permCAF and restCAF subtype. The size of the dots represents the p-value tested by Wilcoxon rank-sum test. **b,** Violin plots comparing the log2 FC of the ratio between the CD8 T cell and Treg fractions in permCAF vs restCAF subtype tumors. Wilcoxon rank-sum test. Correlation of DeCAF permCAF probability in pre-treatment samples with the tumor size change in **c,** permCAF subtype and **d,** restCAF subtype. Spearman correlation. **e, f,** Correlation of the change in neutrophil fraction and M2 macrophages fraction between pre- and post-treatment tumor samples with the percent change in tumor size. Spearman correlation. **g, h,** Boxplot showing the neutrophil and M2 macrophage fractions in pre- and post-treatment samples of permCAF and restCAF subtype tumors, Wilcoxon rank-sum test. **i,** Correlation of permCAF probability between pre- and post-treatment samples (Pearson correlation).

To investigate if DeCAF subtypes are predictive of immunotherapy response in PDAC patients, we examined the Phase 1b trial of FOLFIRINOX in combination with PF-04136309, a CCR2 inhibitor (FFX+PF) which has both pre- and post-treatment samples (Linehan dataset)^27^. Within each subtype, we found that increasing permCAF probability (i.e. increasing permCAFness) in the pre-treatment sample was correlated with a greater percent decrease in tumor size in patients with permCAF (rho = -0.581, p = 0.048, Spearman correlation, Fig. 6c) and restCAF (rho = -0.796, p = 0.002, Spearman correlation, Fig. 6d) subtype tumors. We hypothesized that as CCR2 inhibition targets the recruitment of inflammatory monocytes, our findings may be explained by the enrichment of monocytes in the permCAF subtype samples (Fig. 6a). As this trial had pre- and post-treatment samples, we next looked at the change in neutrophil and M2 macrophages after treatment. We found that decreases in the neutrophil fraction (rho = 0.487, p = 0.016, Spearman correlation) and increases in the M2 macrophage fraction (rho = -0.479, p = 0.018, Spearman correlation) was correlated with tumor response (Fig. 6e,f). In the original trial, CD14+CCR2+ tumor associated macrophages were found to be significantly decreased in FFX+PF treated tumors of 6 patients, but association with response was not avaialble^27^.

We next looked at the relationship between DeCAF subtypes and immune cell population changes in the FFX+PF trial. We found that the decrease in neutrophil and increase in M2 macrophage fractions were specific to permCAF subtype tumors and not found in restCAF subtype tumors (Fig. 6g,h). There was overall correlation of DeCAF score between pre- and post-treatment samples (r = 0.772, p < 0.001, Pearson correlation, Fig. 6i), suggesting that the changes may be less in the CAF subtype, but rather the immune microenvironment associated with the DeCAF subtype. Our results suggest that DeCAF subtypes associate with specific immune microenvironments that may predict immunotherapy response.

### DeCAF subtypes and immunotherapy response in BLCA and cRCC

Clinical trials of immunotherapy in PDAC are limited. As DeCAF subtypes were prognostic in BLCA and cRCC, we evaluated the IMvigor210 trial (NCT02108652) in BLCA^42^ and the IMmotion150 trial (NCT01984242) in cRCC^43^ where patients were treated with the anti-PD-L1 antibody, atezolizumab. In the IMmotion150 trial of metastatic RCC patients, in the atezolizumab only arm, lower DeCAF score or permCAF probability (i.e. increased restCAFness) was numerically, but not significantly associated with having a complete (CR) or partial response (PR) (p = 0.077, t-test, Fig. 7a). In the IMvigor210 trial for metastatic urothelial cancers, we found that patients who had either a CR or PR had significantly lower permCAF probability (i.e. increased restCAFness) (p = 0.014, t-test, Fig. 7b). In addition, patients with permCAF subtype tumors had a mOS of 6.7 months compared to 9.9 months for patients with restCAF tumors (p = 0.043, HR 1.4 [95% CI 1.01, 1.95], Fig. 7c). Finally, similar to PDAC and the TCGA BLCA dataset, in IMvigor210, we found that Ba/Sq subtype tumors were most enriched in the permCAF subtype with 33% (36/109) of Ba/Sq subtype harboring a permCAF subtype (p = 3.81e-05, Fisher’s exact test, Fig. 7d). As expected, higher DeCAF probability was associated with a Ba/Sq tumor subtype as well (p = 5.5e-13, Kruskal-Wallis test, Fig. 7e).

**Fig. 7:**
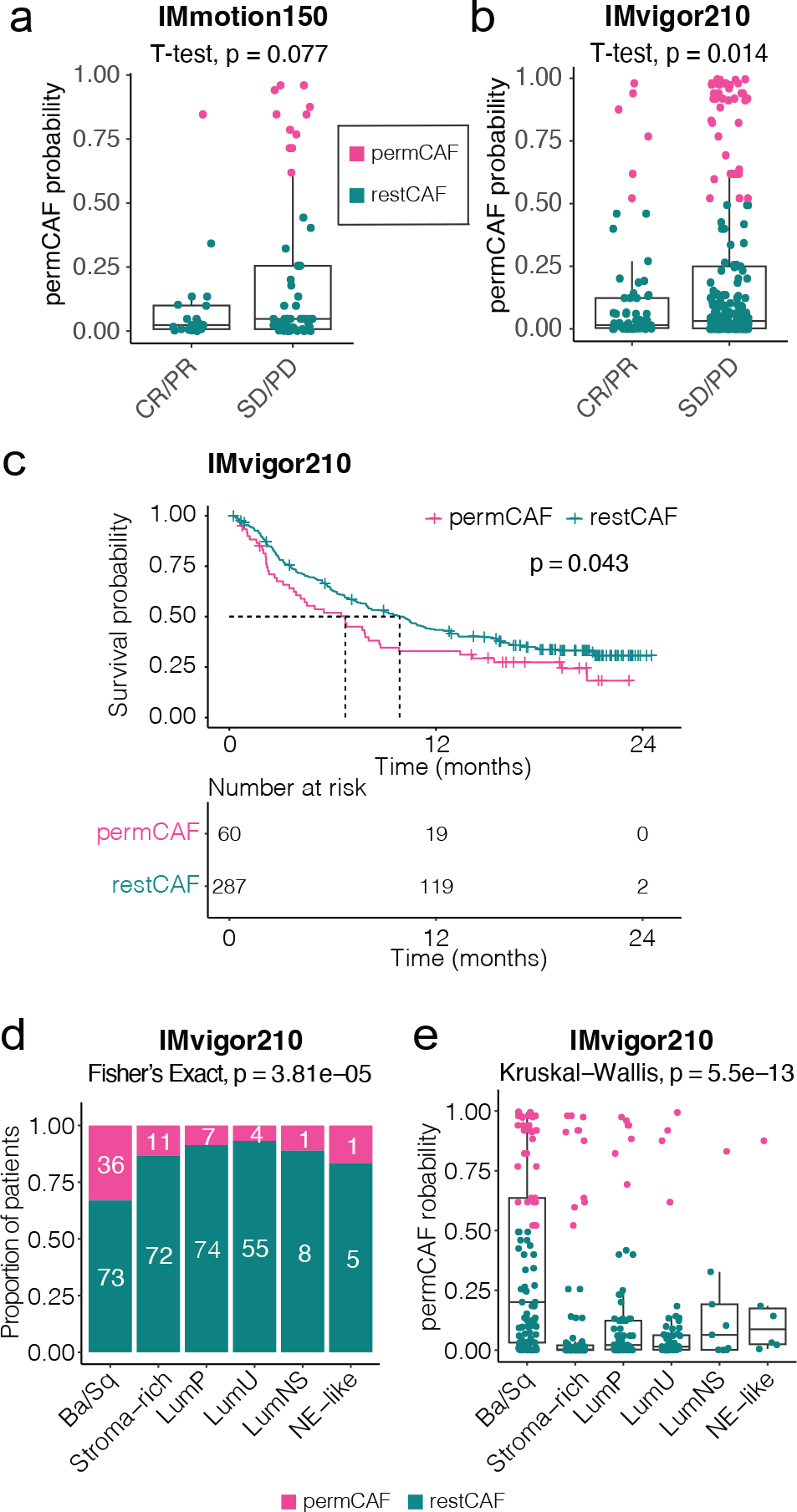
DeCAF subtypes and immunotherapy response in BLCA and RCC. a, b, Boxplot comparing permCAF probability in the patients stratified by ORR in the IMmotion150 (a) and IMvigor 210 trials (b), t-test (CR: complete response, PR: partial response, SD: stable disease, PD: progressive disease). **c,** Kaplan-Meier plot showing OS of patients by DeCAF subtypes in the IMvigor210 trial. **d,** Proportion of patients with permCAF and restCAF subtype tumors within each bladder consensus subtype in IMvigor210, Fisher’s exact test (p <0.001). **e,** Boxplots comparing permCAF probability in patients with different bladder consensus subtype tumors in IMvigor210, Fisher’s exact test (p <0.001, Kruskal-Wallis test).

Therefore, cRCC and bladder cancer patients with restCAF subtype tumors may have an increased overall response rate (ORR) to immune checkpoint inhibition.

## DISCUSSION

Several CAF classifications have been described, but their clinical relevance has been more challenging to translate. Here we leverage the wealth of sc and bulk transcriptomic studies in PDAC to identify the most clinically relevant and robust CAF subtypes. As clustering tends to introduce problems of instability during between-sample normalization when a new sample is added, it cannot be used to call patient subtypes in the clinical setting. For CAF subtype classification, we developed DeCAF, which uses nine pairs of TSP genes to call permCAF vs restCAF subtypes that is robust and replicable. This considerably increases the flexibility and practicality of integrating and studying CAF subtypes in clinical contexts.

A key element of our method includes the utilization of CAF-intrinsic subpopulation marker genes identified using our novel method SCISSORS that precisely identifies CAF clusters and marker genes in scRNAseq data without contamination of other cell types^16–18,20–26^. We show in an independent scRNAseq dataset (UNC-sc) of six PDAC samples that SCISSORS CAF marker genes more precisely discriminate between the respective CAF cell subpopulations without confounding expression in other cell types, compared to previously described CAF subpopulations gene sets^16–18,20–26^. The discriminatory ability of SCISSORS CAF marker genes was critical to successfully deriving and translating CAF-intrinsic subtypes to bulk RNAseq data.

myCAFs were initially described and are thought to be more quiescent with iCAFs as inflammatory and tumor promoting^19^. However, our findings suggest that, in patients, the CAF subpopulations have opposite phenotypes compared to the majority of preclinical studies, with permCAF (myCAF) subtypes with worse prognosis compared to restCAF (iCAF) subtypes. This is in agreement with a recent study that found that myCAFs may be pro-metastatic^44^. Thus, we use the terms permissive and restraining. We find that the histologic features of the stroma in PDAC and mesothelioma have direct associations with the DeCAF subtypes with permCAF subtype tumors having myxoid stroma, and restCAF subtype tumors having fibrous stroma. The DeCAF score allows us to look at mixtures where fibromyxoid (mixed) stroma have intermediate DeCAF scores. Our findings of fibrous histology are consistent with the previously described “deserted subTME” by Grünwald et al. and a “collagen-rich stroma” or “C-stroma” by Ogawa et al. In contrast to prior studies, we do find that the myxoid and fibrous features found on histology also associate with outcome.

Using CIBERSORT, we found the immune landscape of the restCAF subtype to have higher naïve B cell fractions. This is in agreement with previous findings by Grünwald et al., that the deserted subTME trended toward higher B cell marker (CD20) expression in their deep-phenotyping platform for human PDAC tissues^23^. Our findings that permCAF subtype tumors have higher CD8+ T cell percentages is in agreement with prior findings that FAP-stroma is characterized by restricted CD8+ cell infiltrates^24^. The precision of perm-/rest-CAF subtypes in scRNAseq not confounded by immune cells suggests that the CAF subtypes may provide distinct TMEs for different immune cell infiltration.

In agreement with prior studies of CAFs across cancers^35,36^, we found that DeCAF subtypes are prognostic in mesothelioma, cRCC and bladder carcinomas. In bladder cancer, where the PDAC basal-like gene signature can be used to accurately recall the Ba/Sq bladder consensus subtype^16^, both of which have an enrichment of cytokeratins, we find that the permCAF subtype is enriched in both basal bladder Ba/Sq and PDAC basal-like subtype tumors. Furthermore, both tumor and CAF subtype affect prognosis in an additive fashion in bladder cancer patients. Finally, our findings of differential immune microenvironments associated with the permCAF vs restCAF subtype has implications for immunotherapy response in cRCC, bladder and pancreatic cancers. These findings suggest that the biological and therapeutic implications of DeCAF may be extrapolated across these cancer types.

In summary, we find that DeCAF subtypes are histologically distinct, prognostic and predictive of treatment response in multiple cancer types. In pancreatic, cRCC and bladder cancers, the immune microenvironment specific to the subtypes may help predict response to different immunotherapy approaches. Our findings that the biology of previously described iCAF and myCAF subgroups, where iCAFs were thought to be pro-tumorigenic, in patients, is completely reversed, suggests that the interest in targeting iCAF populations may not be as beneficial as originally thought^18^, and may explain some of the disappointing trials to date. We present a clinically tractable CAF subtype SSC, DeCAF, that determines permCAF and restCAF subtypes in patients that may be incorporated into clinical trials, provides a framework for the understanding of CAF subtypes for the development of effective CAF subtype specific therapies, and will facilitate the translation of preclinical studies to patients.

## METHODS

### Marker gene identification in Elyada-sc

SCISSORS includes a carefully designed function, which is a two-step method for the identification of highly cell subpopulation specific genes^26^. Briefly, SCISSORS first derives a candidate gene set by comparing the cell subpopulation of interest to the most related cell subpopulation; then the highly expressed genes from other unrelated cell types are removed from this candidate gene set. In this study using the Elyada-sc dataset, permCAF (myCAF) cells were compared to restCAF (iCAF) and apCAF cells, and the restCAF (iCAF) cells were compared permCAF (myCAF) and apCAF-like cells, to derive a candidate gene set for the cell subpopulation of permCAF (myCAF) and restCAF (iCAF) separately (Wilcoxon rank-sum test, p<0.05, log2 fold change > 0). Then, the permCAF (myCAF) and restCAF (iCAF) candidate gene sets were subjected to a filtering step, in which the highly expressed genes of the non-CAF cells were removed. The highly expressed genes were defined as the top 10% expressed by averaging all the cells within each broad cell cluster. As a result, final gene sets were identified for permCAF (myCAF) and restCAF (iCAF) respectively (Supplementary Table S1).

### Tumor dissociation and library preparation for UNC-sc

Six de-identified primary PDAC samples (UNC-sc, Supplementary Table S2) were collected from the IRB-approved University of North Carolina Lineberger Comprehensive Cancer Center Tissue Procurement Core Facility after IRB exemption in accordance with the U.S. Common Rule. Fresh tissue was dissociated into single cells using Miltenyi human dissociation kits (Miltenyi, 130-095-929) and red blood cells were removed using red blood cell lysis solution (Miltenyi, 130-094-183). Cell counts were performed using an automated cell counter and live cell counts were determined using trypan blue staining. Up to 10,000 cells were encapsulated into droplets for droplet-based 3’ end single-cell RNAseq using Chromium 3’ v3 reagents (10X Genomics). cDNA libraries were quantified using the Qubit dsDNA Assay Kit (Thermo, Q32851) and library quality was assessed with the 4150 Tapestation System (Agilent) and D5000 screen tapes (Agilent, 5067-5588). cDNA libraries were sequenced on a NextSeq500 (Illumina) using NextSeq 500/550 High Output Kit v2.5 (150 Cycles) (Illumina, 20024907) at 200M reads per sample.

### Sample collection and processing for UNC-bulk

129 de-identified primary PDAC patient samples of which 117 have been previously reported (UNC-bulk, Supplementary Table S8) were collected from the IRB-approved University of North Carolina Lineberger Comprehensive Cancer Center Tissue Procurement Core Facility after IRB exemption in accordance with the U.S. Common Rule and were flash frozen in liquid nitrogen. FFPE samples were prepared, hematoxylin and eosin stained. RNA expression libraries were generated for flash frozen samples or FFPE samples, with TruSeq Stranded mRNA kits or with KAPA RNA HyperPrep Kit with RiboErase (HMR) according to the manufacturer’s instructions. Sequencing was performed on the NextSeq500 and NovaSeq6000 Sequencing Systems (Illumina).

### UNC-sc processing

Cell Ranger 6.1.2 was used. BCL files was converted into fastq files using cellranger mkfastq based on bcl2fastq2 (v2.20.0). Fastq files for each sample was then processed by cellranger count to derive unique molecular identifier (UMI) count for each gene, using the human GRCh38 genome. Samples were then aggregated by cellranger aggr.

Aggregated data (filtered_feature_bc_matrix) were analyzed by SCISSORS, which is wrapped around Seurat (v4), for cell clustering. The Quality control steps include 1) the inclusion of genes expressed in more than 2 cells, 2) the inclusion of cells that have the number of genes captured within 200∼2500 (200 < nFeatures < 2500), and 3) the inclusion of cells with mitochondrial reads accounting for less than 5% (percent_MT < 5). The filtered data underwent processing through the PrepareData() function in SCISSORS to obtain the initial clusters with the following parameter settings: n.HVG = 3000, regress.mt = TRUE, n.PC = 20, random.seed = 629, with other parameters using default values. The first-round clusters were annotated using SingleR. Clusters identified as “activated_stellate” were categorized as CAFs. A second-round clustering analysis was performed on the CAF related clusters (0,4,6) using the ReclusterCells function in SCISSORS, with the following parameter settings: merge.clusters = TRUE, use.sct = TRUE, n.HVG = 3000, regress.mt = TRUE, n.PC = 20, resolution.vals = 0.2, k.vals = 57, with other parameters using default values.

### UNC-bulk processing

Raw base call (BCL) files were converted to fastq files using bcl2fastq2 (v2.19.0). RefSeq assembly (GCF_000001405.40) of the human reference genome GRCh38.p14 was used as the reference for gene quantification by Salmon 1.9.0^45^ (“-- gcBias -- seqBias”). The total expected read counts per gene were normalized to transcripts per million (TPM).

### Public bulk datasets and sample inclusion

Eleven bulk transcriptomics datasets were obtained from public sources (Supplementary Table S4). Gene expression quantifications was used ‘as-is’ with respect to the original publications when possible, i.e. data were not re-aligned or re-quantified; gene-level expression estimates were used either in the unit of TPM (transcripts per million) or FPKM (fragments per kilobase per million reads), depending on the study. When preprocessed gene expression data were not available, for training datasets, the most similar methods were used to process the data; for validation and independent datasets, Salmon 1.9.0^45^ using RefSeq assembly (GCF_000001405.40) of the human reference genome GRCh38.p14 was used to derive gene expression levels (Supplementary Table S4). Samples from the public datasets were filtered to include only non-metastatic primary PDAC samples (Supplementary Table S5). For the Grünwald and Olive datasets, which are microdissected, only stroma samples were included.

Bladder cancer subtyping calls were made on log2 transformed upper-quartile normalized expression data using the BLCA subtyping and consensusMIBC R package^46^. Within the UROMOL and IMvigor210 datasets, all samples with gene expression data were used in the analysis. For the TCGA_BLCA cohort, only tumors from patients with stage M0 disease were included in the analysis.

### Consensus clustering (CC)

SCISSORS permCAF and restCAF genes were derived as described above and ranked by fold enrichment to generate the top25 genes for each subtype. Gene sets of the Moffitt stroma, Elyada and Maurer schemas were collected from each study respectively. For each of the subtyping schemas in each of the 11 public transcriptomics datasets, unsupervised CC was applied using the ConsensusClusterPlus (version 1.56.0) package in R for genes (rows) and samples (columns) separately. Data matrices were subjected to log2 transformation and column-wise quantile normalization. Note that the data were not normalized row-wise, as that may force the clusters to have similar sizes, instead of deriving the reflective number of patients in each cluster. For clustering of samples, a distance matrix was derived based on Pearson correlation. Then CC was applied to this distance matrix to derive two sample clusters (K = 2), which consisted of 1,000 iterations of k-means clustering using Euclidean distance, with 80% items hold-out at each iteration. The number of K was determined empirically by visual inspection to derive clusters of samples that were most representative of the CAF subtypes. For clustering of genes, a distance matrix was derived based on Pearson correlation. Then CC was applied to this distance matrix derive two gene clusters (K = 2), which consisted of 200 iterations of k-means clustering using Euclidean distance, with 80% items hold-out at each iteration.

### Generation of CC labels for classifier training and validation

To derive confident labels for classifier training and validation, the CC method mentioned above were adapted. Specifically, the CC starts at sample-wise K=2, with Ks increasing step by step for inspection of clustering performance and sample-gene associations. The resultant clusters were then labeled as “permCAF”, “Mixed permCAF”, “Mixed”, “Mixed restCAF”, “restCAF” and “Absent” based on gene expressions (Supplementary Figure S3). A dataset does not necessarily have every one of the 6 cluster categories. “Mixed permCAF” was then merged with ”permCAF”, and “Mixed restCAF” merged with “restCAF”. These merged “permCAF” and “restCAF” labels were used for training and validation of DeCAF.

### DeCAF classifier training

#### Candidate gene ranking

SCISSORS CAF genes were ranked based on the consistency of their differential expression (DE) statistics between CC-based subtypes in each individual training dataset (Supplementary Table S5). A cross-study DE consistency score was obtained by summing the -log10 p-values (Wilcoxon rank-sum test) and ranking them from high to low. The top 25% of this set (consistent DE genes) were considered for model training and the genes where the direction of up- regulation or down-regulation of them were not consistent in the subtypes were removed. The remaining genes then formed our final candidate gene set for downstream steps.

#### Rationale of using kTSP for binary classification

Let us define a gene pair (*g*_*dis*_,*g*_*dit*_), where *g*_*dis*_ is the expression of gene *s* for subject *i* in study *d*, and *g*_*dit*_ is the expression of gene *t* for the same subject and study. A TSP is an indicator variable based on this gene pair, 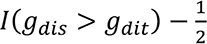, where the value represents which gene in the pair has higher expression in subject *i* from study 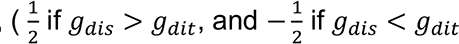 otherwise). The TSP method was originally proposed in the context of binary classification [cite]In traditional applications, a single TSP (k = 1) is selected out of the set of all possible gene pairs, in which case, 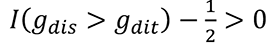 implies subtype A with high probability, otherwise subtype B is implied [cite]. We view such binary variables as “biological switches” indicating how pairs of genes are expressed relative to clinical outcome.

In the kTSP setting, class prediction reduces to verifying whether the sum across k selected TSPs is greater than 0:

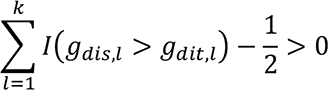

This reduces to a majority vote across the selected k TSPs, where the contribution of each of the k TSPs are equally weighted to select subtype A if the above sum is greater than 0, and subtype B otherwise. However, some TSPs may be more informative than others, so we utilized penalized logistic regression [cite] to jointly estimate the effect of each of the k selected TSPs in predicting binary subtype, and to further remove TSPs with weak or redundant effects. Predicted probabilities of permCAF subtype membership (DeCAF score) may then be obtained from the fitted logistic regression model on our training samples, where values greater than 0.5 indicate predicted membership to the permCAF subtype and restCAF otherwise.

#### Horizontal data integration and kTSP selection via switchBox

To apply the top scoring pairs transformation, we utilize the switchBox R package (version 1.28.0) [cite] to enumerate all possible gene pairs based on our final candidate gene list and training samples (function SWAP.KTSP.Train, with optimal parameters featureNo=100, krange = 50, FilterFunc = NULL). Given the large number of potential gene pairs based on this list, in addition to the strong correlation between gene pairs sharing the same genes, the switchBox package utilizes a greedy algorithm to select from this list a subset of gene pairs that are helpful for prediction, given the set of training labels. We merge data from each training dataset without normalization prior to applying switchBox, as the method only looks at the relative gene expression ranking within each sample from each study. The method then selects a subset of k TSPs, where k is determined through a greedy optimization procedure.

#### Model training based on selected kTSPs

To remove redundant TSPs and to jointly estimate their contribution in predicting subtype in our training samples, we utilize the ncvreg R package (version 3.13.0) to fit a penalized logistic regression model based upon the selected TSPs from switchBox. Our design matrix is an N x (k+1) matrix, where the first column pertains to the intercept and the remaining k columns pertains to the k selected TSPs from switchBox. Here N is the total number of training samples (Supplementary Table S4). Each TSP in the design matrix is represented as a binary vector, taking on the value of 1 if gene A’s expression is greater than gene B’s expression. Our outcome variable here is binary subtype (1 = permCAF, 0 otherwise). We utilize optional parameters alpha = 0.05 and nfolds = N. We allow for correlation between TSPs by setting the ncvreg alpha parameter to 0.05 in order to shrink the coefficients of highly correlated TSPs and also remove correlated uninformative TSPs from the model. We set nfolds = N to apply leave one out cross validation in order to choose the optimal MCP penalty tuning parameter for variable selection, where the optimal tuning parameter is the one that minimizes the cross-validation error of the fitted model. Our final model then reports the set of coefficients estimated for each of the kTSPs, where each coefficient may be interpreted as the change in log odds of a patient being part of the permCAF subtype when the lth TSP is equal to 1, given the others in the model. TSPs with coefficient of 0 are those that have been removed from the model for either weak effect or redundancy with other TSPs.

### Final kTSP model

As illustrated in Figure 3A, 9 pairs of kTSP genes were evaluated for their relative ranking in a new patient. A value of 1 is assigned if the permCAF gene in a TSP has greater expression than the restCAF gene in that patient (and 0 assigned other wise) creating *X*_*i,new*_, a 1 x (k +1) TSP predictor vector. These values are then multiplied by the corresponding set of estimated TSP model coefficients, *β̂*, obtained from the fitted penalized logistic regression model. The intercept term is included to correct for estimated baseline effects. These values are summed to get the patient “DeCAF Score” *X*_*i,new*_. This score is then converted to a predicted probability of belonging to the permCAF subtype by computing its inverse logit:

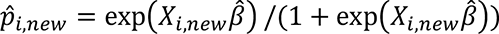

Values greater than or equal to 0.5 indicated predicted permCAF subtype membership, and those less than 0.5 are predicted to be of the restCAF subtype. This is equivalent to determining whether *X*_*i,new*_*β̂* > 0 (permCAF subtype) vs *X*_*i,new*_*β̂* < 0 (restCAF subtype). Thus, the DeCAF score, *X*_*i,new*_*β̂*, may also be utilized as a continuous score for classification. Therefore, prediction in new samples, such as from our validation datasets, reduces to simply checking the relative expression of each gene within the set of TSPs.

For all discussions regarding classifier performance, we obtain the predicted subtypes in the manner described above. The level of confidence in the prediction can be determined based upon the distance of *X*_*i,new*_*β̂* from 0.5, where values closer to 0.5 indicate lower confidence in the predicted subtype and higher confidence otherwise.

### Validation of DeCAF

The performance of the final DeCAF model in the training set was measured using leave-one-out cross validation. This was implemented using the cv.ncvreg function from the ncvreg (version 3.13.0). Since the outcome is binary, the cross-validation error is measured via the leave-one-out cross-validated deviance of the logistic regression model.

Next, the performance of the DeCAF model was evaluated in the validation sets. The validation set is composed of seven separate studies. To account for the variability between studies, a fully non-parametric ROC curve was used for this meta-analysis. The ROC curve was constructed and the inter-study variability was measured using the study random-effects model^47^ and implemented in the metaROC function from the nsROC R package (version 1.1). All validation metrics compared the DeCAF classifier to the combined “Mixed permCAF” and “permCAF” calls and combined “Mixed restCAF” and “restCAF” clustering calls.

### Survival analysis

For pooled survival analysis, patients with subtype calls, OS time and event were involved. For the Linehan dataset, where patients received treatments, only pre-treatment samples were included when both pre-treatment and post-treatment samples were available. For samples that are duplicated in the PACA_AU_seq and PACA_AU_array datasets, only the sample from PACA_AU_seq was used for survival analysis.

Overall survival estimates were calculated using the Kaplan-Meier method. Association between overall survival and individual covariates such as subtype were evaluated via the cox proportional hazards models using the coxph function from the ‘survival’ R package (version 3.2-13), where a given subtyping schema was considered as a multi-level categorical predictor. The log-rank test was used to evaluate overall association of a subtyping schema with overall survival and derive the p-values. In the pooled analyses, a stratified cox proportional hazards model was utilized, where dataset of origin was used as a stratification factor to account for variation in baseline hazard across studies.

### CIBERSORT

The analytical tool CIBERSORT^48^ was employed to estimate the percentage of cell types for each sample of 12 bulk transcriptomics data. This estimation was performed using the LM22 signature matrix, which encompasses 547 genes that distinguish 22 human hematopoietic cell phenotypes. With the percentage results obtained from CIBERSORT, we compared the results across two types of DeCAF calls for the samples in each dataset. The Wilcoxon rank-sum test was employed to assess whether there were significant differences. The log2 fold change of the median was utilized to quantify the magnitude of the observed differences.

### Data Availability

Bulk public datasets downloaded and involved in this study were summarized in Supplementary Table S4. UNC-sc and UNC-bulk data will be uploaded to public repository upon acceptance of the manuscript and are currently available upon request by the reviewers.

### Code Availability

The DeCAF classifier was deposited as a GitHub repository at: https://github.com/jjyeh-unc/decaf. All scripts involved in generating results and figures are available upon request.

### Authors’ Disclosures

A patent application is planned, but not yet filed for this work: Authors/inventors: Jen Jen Yeh, Naim U Rashid, Elena V Kharitonova and Xianlu L Peng by the University of North Carolina Chapel Hill. PurIST, used in the manuscript, has a patent pending 17/601,002 by the University of North Carolina Chapel Hill. Inventors who are authors: Xianlu L Peng, Jen Jen Yeh, Naim U Rashid. PurIST was licensed to GeneCentric Therapeutics, Inc. which had no participation in or knowledge of the current work.

### Disclaimer

The study design, data collection and analysis, decision to publish, and manuscript preparation were conducted independently of any involvement from the funders.

## Acknowledgments

We would like to thank the University of North Carolina at Chapel Hill and the Research Computing group for providing computational resources and support. We thank the UNC Lineberger Comprehensive Cancer Center Tissue Procurement Facility and Preclinical Studies Core for excellent support and assistance. We thank Barbara Grünwald and members of the NCI Pancreatic Stroma Research Consortium for in depth discussions and insights.

## Notes

### Competing Interest Statement

The authors have declared no competing interest.

