## Supplementary Fig. 1 for "Determination of permissive and restraining cancer-associated fibroblast (DeCAF) subtypes"

**Supplementary Fig. 1. Subtyping PDAC patient samples by consensus clustering.** Heatmaps showing consensus clustering (CC) of the datasets using SCISSORS genes.

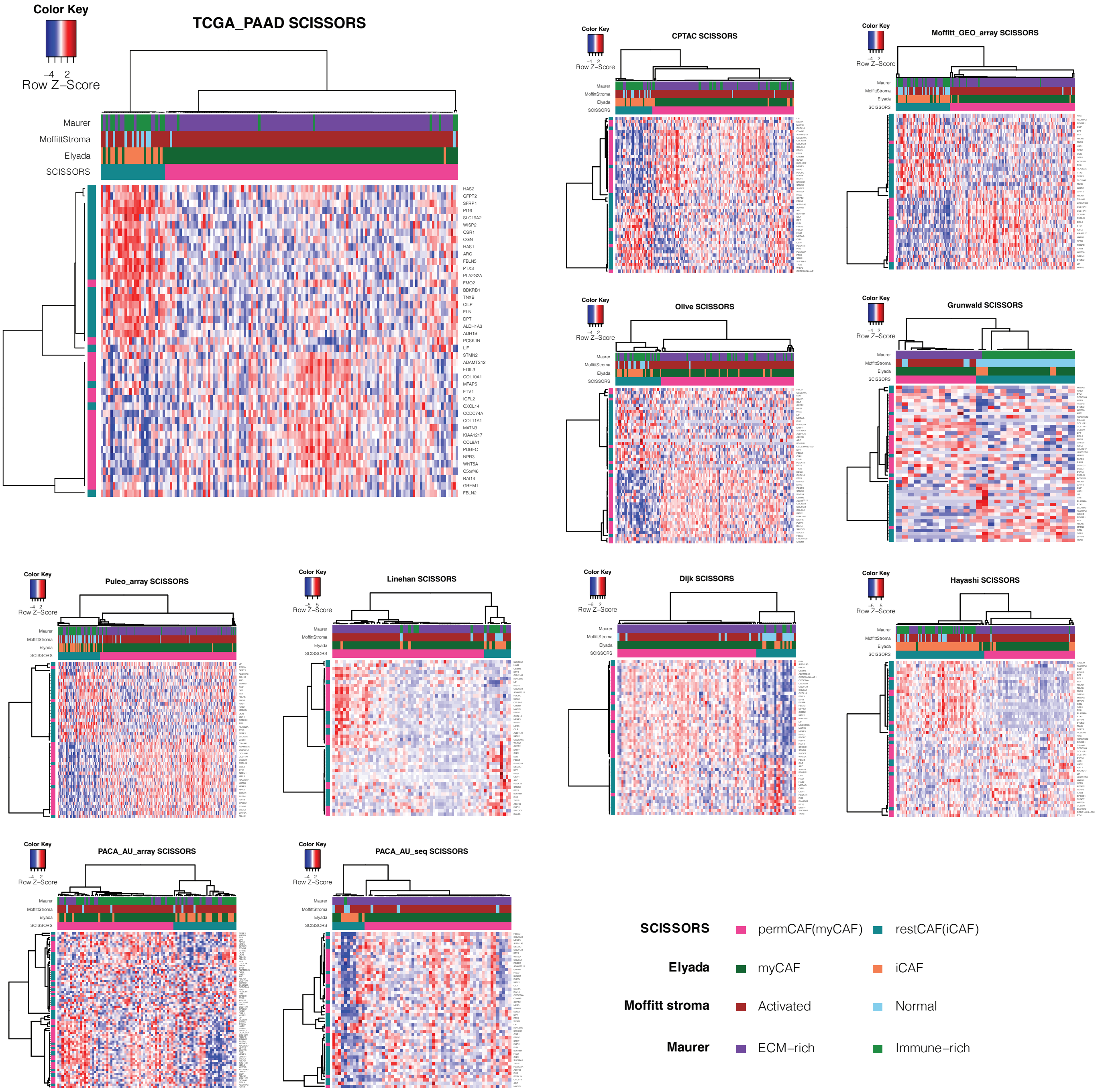
