## Supplementary Fig. 2 for "Determination of permissive and restraining cancer-associated fibroblast (DeCAF) subtypes"

**Supplementary Fig.2: Assigning gold standard CAF subtype labels using consensus clustering (CC).** Heatmaps showing CC of the datasets using SCISSORS CAF genes to derive training and validation labels. CC used at variable resolution of Ks (vK) to label each sample as one subtype of permCAF, mixed permCAF, mixed, mixed restCAF, restCAF or absent.

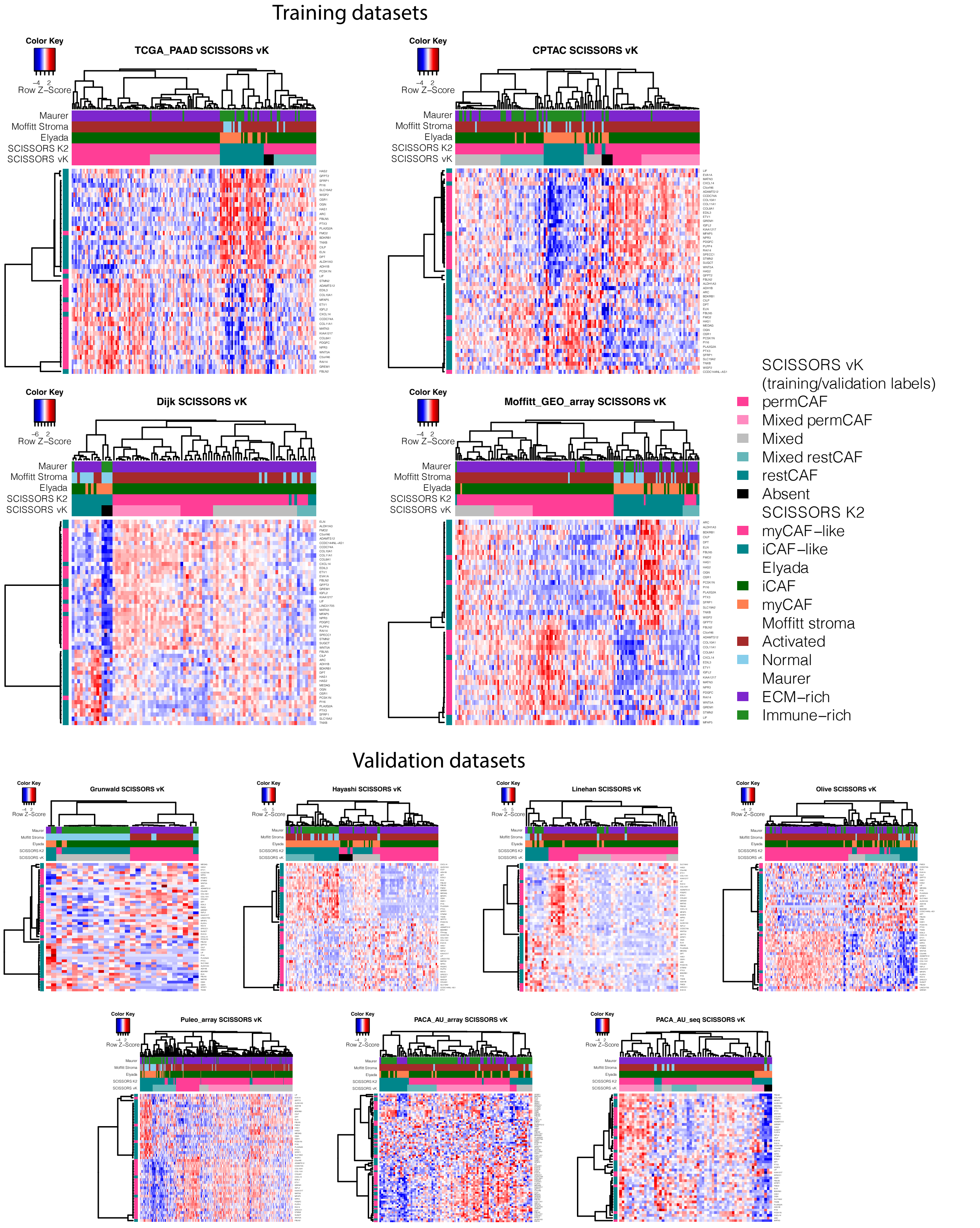
