## Supplementary Fig. 3 for "Determination of permissive and restraining cancer-associated fibroblast (DeCAF) subtypes"

**Supplementary Fig. 3: Number of overlappings between different CAF marker sets.** The intensity of the color on the heatmap showing the absolute number of gene overlaps, with greater redness indicating a larger number.

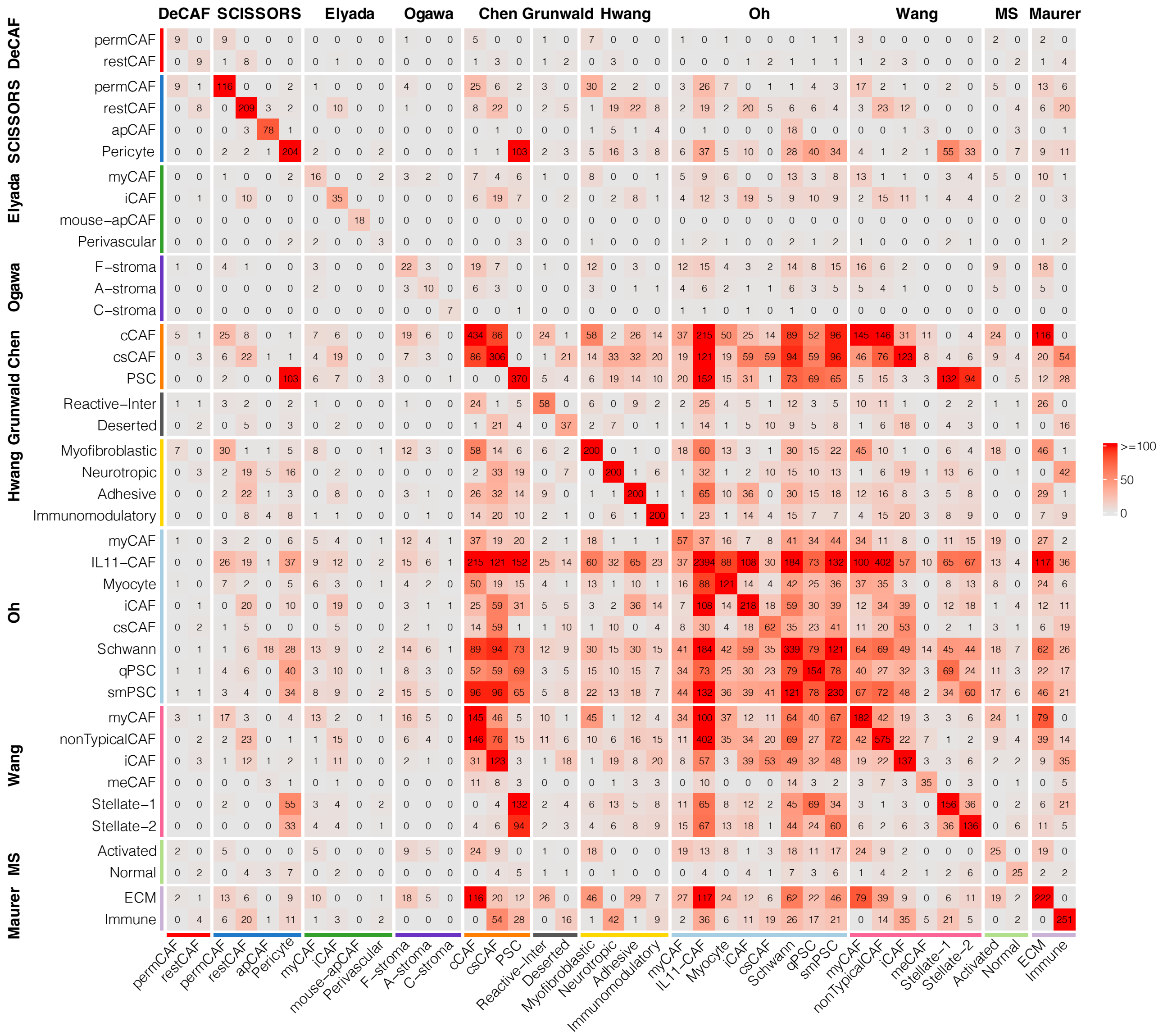
